# Differential protein phosphorylation affects the localisation of two secreted *Toxoplasma* proteins and is widespread during stage conversion

**DOI:** 10.1101/2020.04.09.035279

**Authors:** Joanna Young, Malgorzata Broncel, Helena Teague, Matt Russell, Olivia McGovern, Matt Renshaw, David Frith, Bram Snijders, Lucy Collinson, Vern Carruthers, Sarah Ewald, Moritz Treeck

**Affiliations:** Signalling in Apicomplexan Parasites Laboratory, The Francis Crick Institute, London, UK; Electron Microscopy Science Technology Platform, The Francis Crick Institute, London, UK; Department of Microbiology and Immunology, University of Michigan Medical School, Ann Arbor, Michigan, USA; Advanced Light Microscopy Science Technology Platform, The Francis Crick Institute, London, UK; Proteomics Science Technology Platform, The Francis Crick Institute, London, UK; Department of Microbiology, Immunology and Cancer Biology and the Carter Immunology Center, University of Virginia, Charlottesville, Virginia, USA

**Keywords:** Host-Pathogen Interaction, virulence factors, phosphorylation, chronic infection

## Abstract

The intracellular parasite *Toxoplasma gondii* resides within a membrane bound parasitophorous vacuole (PV) and secretes an array of proteins to establish this replicative niche. It has been shown previously that *Toxoplasma* both secretes kinases and that numerous proteins are phosphorylated after secretion. Here we assess the role of phosphorylation of SFP1 and the related GRA29, two secreted proteins with unknown function. We show that both proteins form stranded structures in the PV that are independent of the previously described intravacuolar network or actin. GRA29 likely acts as a seed for SFP1 strand formation, and these structures can form independently of other *Toxoplasma* secreted proteins. We show that an unstructured region at the C-terminus of SFP1 and GRA29 is required for the formation of strands and that mimicking phosphorylation of this domain negatively regulates strand development. When tachyzoites convert to chronic stage bradyzoites, both proteins show a dispersed localisation throughout the cyst matrix. Many secreted proteins are reported to dynamically redistribute as the cyst forms and secreted kinases are known to play a role in cyst formation. Using quantitative phosphoproteome and proteome analysis comparing tachyzoite and early bradyzoite stages, we reveal widespread differential phosphorylation of secreted proteins. These data support a model in which secreted kinases and phosphatases are important to dynamically regulate parasite secreted proteins during stage conversion.

**IMPORTANCE:** *Toxoplasma gondii* is a common parasite that infects up to one third of the human population. Initially the parasite grows rapidly, infecting and destroying cells of the host, but subsequently switches to a slow-growing form and establishes chronic infection. In both stages the parasite lives within a membrane bound vacuole within the host cell, but in the chronic stage a durable cyst wall is synthesized that provides protection to the parasite during transmission to a new host. *Toxoplasma* secretes proteins into the vacuole to build its replicative niche and previous studies identified many of these proteins as phosphorylated. We investigate two secreted proteins and show that phosphorylation plays an important role in their regulation. We also observed widespread phosphorylation of secreted proteins when parasites convert from acute to chronic stages, providing new insight into how the cyst wall may be dynamically regulated.

## INTRODUCTION

The apicomplexan parasite *Toxoplasma gondii* is widespread infecting approximately one third of the world’s population (1). Although *Toxoplasma* infection is predominantly asymptomatic in healthy hosts, complications occur in the immunocompromised, such as cancer and HIV patients, and in pregnant women. In these patients, infection can cause encephalitis, or severely damage the unborn foetus during pregnancy (2). Additionally, *Toxoplasma* is a leading cause of retinocharditis (3), and an expansion of strain diversity in South America is causing substantial ocular disease (4).

*Toxoplasma* actively invades nucleated cells of virtually any warm-blooded animal and subsequently replicates within a membrane-bound parasitophorous vacuole (PV). During the acute stage of infection, the rapidly growing tachyzoites complete rounds of invasion and lysis of host cells and disseminate around the body. Upon immune pressure or other stress, the parasite converts to slow-growing bradyzoites that form cysts, predominantly in skeletal muscle and the brain (5). The cyst is a modified PV with a protective cyst wall that allows the parasite to establish chronic, perhaps lifelong infection of the host.

During infection *Toxoplasma* secretes proteins that subvert host cell signalling and develops its replicative niche within the PV (6, 7). The repertoire of secreted proteins is thought to include up to 200 proteins (ToxoDB, LOPIT dataset) including 50 kinases/pseudokinases (8), although the functions of many of these remain unknown. These proteins are secreted from the rhoptries or dense granules and are called ROPs or GRAs, respectively. Secreted proteins extensively modify the PV allowing selective passage of molecules and proteins across the PV membrane. For example, GRA17 and GRA23 form pores in the PV membrane allowing the passage of small molecules into the PV (9), while the MYR complex mediates the transport of secreted proteins to the host cell (10, 11). Additionally, the parasite develops a complex set of membrane structures within the PV including GRA7-lined invaginations (12) and a network of tubules called the intravacuolar network (IVN) (13). The formation of this network is dependent on GRA2 and the accessory protein GRA6, and is required for full virulence in mice (14, 15). Both structures have been reported to play a role in scavenging nutrients from the host cell with the uptake of host cell proteins (16), lipid droplets (17), and endolysosomal compartments (18).

Previous phosphoproteome analysis revealed that a large number of *Toxoplasma* secreted proteins are phosphorylated after their release (19), raising the intriguing possibility that their function is dynamically regulated. Indeed, the PV localised kinase WNG1 was shown to contribute to formation of a functional IVN (20). Furthermore, it was recently shown that IVN associated GRAs show dynamic localisation patterns as the parasite remodels the PV to form the chronic stage cyst (21). Here we analyse two secreted, strand forming proteins that are phosphorylated after secretion and show that phosphorylation regulates their localisation, disrupting normal strand formation. Both proteins disperse in chronic stage cysts, so we expand the analysis to compare the phosphoproteome of acute and chronic stage parasites. We show that many GRAs are differentially phosphorylated between these stages, suggesting that phosphorylation of secreted proteins may be a key determinant for dynamic restructuring of the replicative niche of *Toxoplasma*.

## RESULTS

### A secreted protein forms strand-like structures in the parasitophorous vacuole independently of the intravacuolar network and actin

To investigate the role of phosphorylation of secreted proteins we selected TGGT1_289540 as an ideal candidate. It is substantially phosphorylated after secretion (19) and contains a localised cluster of phosphorylation sites in its C-terminus, allowing for targeted genetic mutagenesis. TGGT1_289540 contains a signal peptide, four coiled-coil domains and an unstructured C-terminus which contains 8 phosphorylation sites (Fig. 1A). Three further phosphorylation sites are located within the rest of the protein. To first verify that TGGT1_289540 is a secreted protein, we expressed a myc-tagged version of the protein which localised to the PV. Interestingly, in contrast to most GRAs that are either found to fill the space between parasites, or are found associated with the PV membrane, TGGT1_289540 appeared in strand- or filament-like structures (Fig.1B). Western-blotting confirmed the presence of a single isoform of myc-tagged protein at the predicted size of ^~^100kDa in transgenic parasites, but not in a WT control (Fig. 1C).

**Figure 1.**
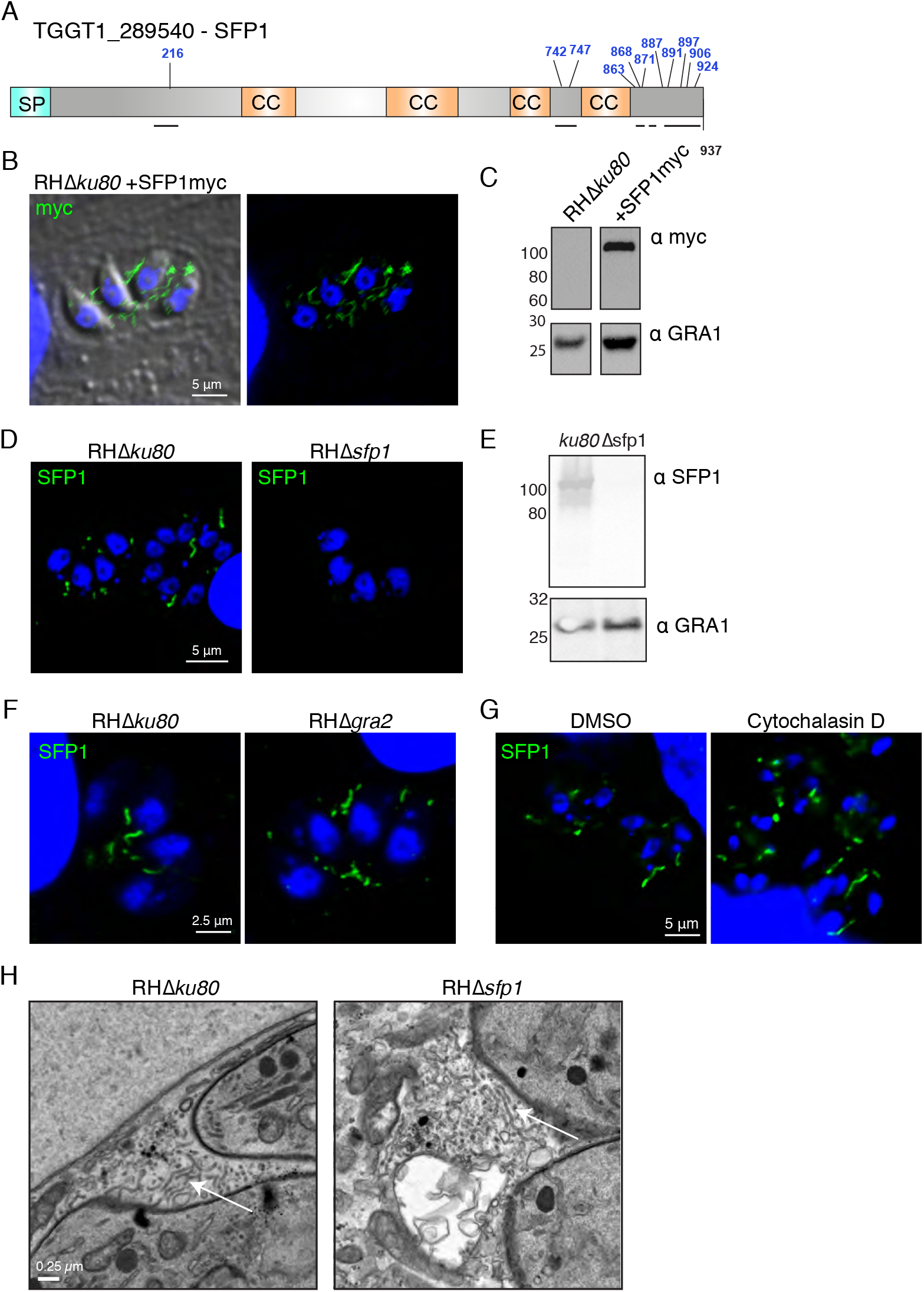
SFP1 forms strands in the parasitophorous vacuole that are distinct from the intravacuolar network. **a)** Schematic of TGGT1_289540 (SFP1) indicating predicted domains and previously identified phosphosites (shown in blue). Signal peptide (SP), coiled coil (CC). Black lines show predicted unstructured regions. **b)** Immunofluorescence images of HFFs infected with *RHDku80* parasites expressing SFP1-myc. Anti-myc antibody (green) and DAPI (blue). **c)** Western blot of immunoprecipitated TGGT1_289540::myc. Anti-GRA1 antibodies were used to probe the non-bounds fraction as a loading control. **d)** Immunofluorescence images of HFFs infected with *RHDku80* and *RHDsfp1* parasites. Polyclonal anti-SFP1 antibodies (green) and DAPI (blue). **e)** Western blot analysis of lysates from HFFs infected with RHΔ*ku80* and RHΔ*sfp1* parasites probed with anti-SFP1 and anti-GRA1 antibodies. **f)** SFP1 strands do not depend on GRA2 and the IVN. Immunofluorescence analysis of RHΔ*ku80* and RHΔ*gra2* infected HFFs. Anti-SFP1 antibodies (green) and DAPI (blue). **g)** SFP1 strands do not depend on actin filamentation. Immunofluorescence analysis of infected HFFs treated with 1uM Cytochalasin D or DMSO control. Anti-SFP1 antibody (green) and DAPI (blue). **h)** SFP1 is not required for IVN formation. Electron micrographs of HFFs infected with RHΔ*ku80* and RHΔ*sfp1* for 16 h. Examples of Intravacuolar network tubules are indicated by arrows.

To verify this unusual distribution of TGGT1_289540 in the PV, we generated polyclonal antibodies against the recombinant protein. The antibody recognised a band in Western blot that was absent in WB and IFA when the *TGGT1_289540* locus was disrupted (Fig. 1D,E). As was the case with the myc-tagged isoform, analysis of the endogenous protein showed a filamentous distribution in the PV (Fig. 1D).

As the *Toxoplasma* PV contains a distinct IVN of membranous tubules we tested whether the strands were related to these structures. The formation of the IVN is dependent on GRA2 so we assessed TGGT1_289540 localisation in RHΔ*gra2* parasites that lack the IVN (14). No obvious difference in filamentous TGGT1_289540 localisation was observed (Fig. 1F), indicating that the filaments are distinct from, and their formation does not depend on the IVN. We therefore named the protein Strand Forming Protein 1 (SFP1 hereafter).

We hypothesised that SFP1 strands could function in transport between the parasites and the host cell. *Toxoplasma* endocytoses proteins from the host cell through an unknown mechanism (16) so we assessed whether SFP1 plays a role in the uptake of proteins from the host cell into the parasite at 3h post infection. As endocytosed material is rapidly digested, this can only be observed upon inhibition of cathepsin protease L function with the inhibitor morpholinurea-leucine-homophenylalanine-vinyl phenyl sulphone (LHVS)(16). Under these conditions there was no difference in the proportion of parasites that had taken up Venus fluorescent protein from the host cell indicating SFP1 does not contribute to protein uptake (Fig. S1A). *Toxoplasma* upregulates host c-Myc in infected cells, a phenotype that is dependent on protein translocation from the PV into the host cell (10, 22). To test whether SFP1 plays a role in protein export into the host cell we compared the ability of wild type and Δ*sfp1* parasites to upregulate host c-Myc. No differences where observed in host cell c-Myc upregulation, indicating that SFP1 does not play a role in protein export from the PV (Fig. S1B).

It was recently shown that long filaments of actin extend into the PV during *Toxoplasma* infection (23). We therefore tested whether treatment with cytochalasin D, to depolymerise actin, would disrupt SFP1 strand formation. We still saw substantial filamentation of SFP1 in the presence of cytochalasin D, indicating that actin is not important in this process (Fig. 1G).

While the presence of the IVN is not important for SFP1 filament formation, the opposite could be true, that is, that SFP1 may be important for the formation of the IVN. However, IVN formation was not affected in the absence of SFP1 as shown by transmission electron microscopy (Fig. 1H). No obvious abnormalities in the overall PV organisation or PVM structure were observed in RHΔ*sfp1* parasites.

Collectively these data indicate that SFP1, a protein that was reported to be phosphorylated after secretion, localises to the PV, and forms novel, IVN- and actin-independent filamentous structures. These, as suggested by our data, do not contribute to protein uptake from, or protein export into the host cell.

### A second SFP1 related protein, GRA29, forms related structures

As secreted protein families can form by gene duplication and diversification, we looked for potential paralogs of SFP1 in the *Toxoplasma* genome using BlastP. We identified a single protein (TGGT1_269690, GRA29) with 24 % identity and a similar architecture (Fig. 2A & S2) of a signal peptide, coiled-coil domains and, like SFP1, C terminal phosphorylation sites that appear phosphorylated after secretion (19). To compare localisation of this protein with that of SFP1 we generated an HA-tagged version which by IFA localised to the PV and Western blot showed an expected size of ^~^94 kDa (Fig. 2B,C). In contrast to SFP1 however, we saw smaller filamentous structures and also large puncta associated with the parasites (Fig. 2B) consistent with the localisation reported by Nadiparum *et al(24)*. To investigate whether SFP1 and GRA29 were within the same structures, we performed super-resolution microscopy using anti-SFP1 antibodies and anti-HA antibodies to localise GRA29. At this improved resolution GRA29 puncta appeared as round spheres or doughnuts, often with SFP1 filaments associated or branching out from them (Fig. 2D). We also observed filamentous SFP1 with smaller dots and strands of GRA29. To further investigate these unusual sphere-like structures we performed correlative light-electron microscopy (CLEM) to identify the subcellular structures. This revealed that GRA29::HA accumulates in electron dense particles in the PV that appear to contain no membrane and resembles proteinaceous aggregates (Figure S3).

**Figure 2.**
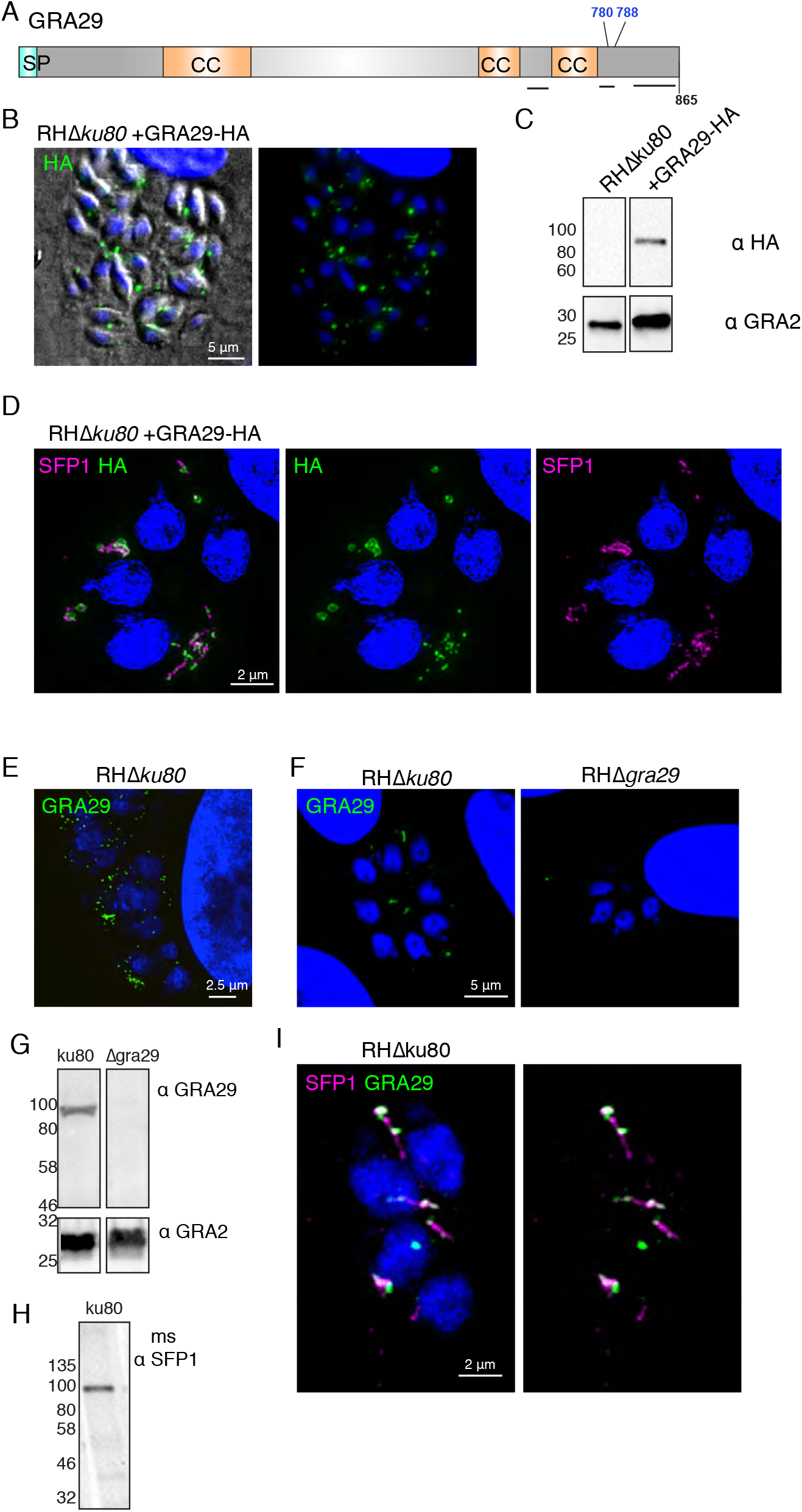
GRA29 shares homology with SFP1 and forms similar structures in the PV. **a)** Schematic of GRA29 indicating predicted domains and previously identified phosphosites (shown in blue). Signal peptide (SP), coiled coil (CC). Black lines show predicted unstructured regions. **b)** Immunofluorescence analysis of HFFs infected with RHΔ*ku80*::GRA29-HA. Anti-HA antibodies (green) and DAPI (blue). **c)** Western blot analysis of RHΔ*ku80* or RHΔ*ku80*::GRA29-HA infected HFFs with anti-HA and anti-GRA2 antibodies to verify expression of HA-tagged protein. **d)** Structured Illumination microscopy (SIM) of HFFs infected with RHΔ*ku80*::GRA29-HA. Anti-HA (green), anti-SFP1 (magenta) and DAPI (blue). **e)** SIM analysis of RHΔku80. Anti-GRA29 antibodies (green) and DAPI (blue) **f)** Immunofluorescence analysis of HFFs infected with RHΔ*ku80* and RHΔ*gra29* parasites. Anti-GRA29 (green) and DAPI (blue). **g)** Western blot of RHΔ*ku80* and RHΔ*gra29* infected HFFs probed with anti-GRA29 and anti-GRA2 antibodies. **h)** Western blot of RHΔ*ku80* infected HFFs probed with mouse anti-SFP1 antibodies generated against SFP1 C-terminal showing a band of the expected size. **i)** SIM analysis of RHΔ*ku80* infected HFFs. Mouse anti-SFP1 (magenta), anti-GRA29 (green) and DAPI (blue). See also Fig. S3.

To further investigate and verify GRA29::HA structures, we raised polyclonal antibodies against recombinant GRA29. With the antibodies against endogenous protein we observed only short strands and small puncta in the PV that did not resemble the HA-tagged parasite line (Fig. 2E,F). The specificity of the antibodies was verified by Western blot and IFA, which showed the expected molecular weight and positive IFA signal in WT, but not in GRA29 KO parasites (Fig. 2F,G). This indicates that the addition of the C-terminal HA tag disrupts GRA29 localisation, inducing protein aggregates, suggesting a potentially important role of the C-terminal region. In order to colocalise endogenous non-tagged SFP1 and GRA29 we generated antibodies against SFP1 peptides in mice which showed the expected reactivity in IFA and Western blot (Fig. 2H, I). Co-localisation of SFP1 and GRA29 revealed that both appear in similar structures, with GRA29 often present at either end of SFP1 filaments (Fig. 2I).

Collectively these data show that SFP1 and GRA29 form novel filamentous structures in the PV. C-terminal tagging of GRA29 alters its localisation, either causing or stabilising unusual structures that SFP1 can still associate with.

### GRA29 initiates SFP1 strand formation, even in the absence of other *Toxoplasma* proteins

As SFP1 and GRA29 appeared to localise within the same structures we hypothesised that they may show an interdependence. To address this, we localised SFP1 in RHΔ*gra29*, and GRA29 in RHΔ*sfp1* parasite lines using the specific antibodies raised. Whereas no alteration in GRA29 was observed in RHΔ*sfp1*, SFP1 formed fewer and longer filaments in the absence of GRA29 (Fig. 3A). To identify whether any other *Toxoplasma* proteins are required for i) SFP1 filament formation and ii) SFP1 filament initiation, we expressed both SFP1 and GRA29, lacking their respective signal peptides, in human foreskin fibroblasts (HFFs). Both, SFP1 and GRA29 formed long filamentous structures in HFFs (Fig. 3B), often filling the complete cytoplasm, and in the case of GRA29, sometimes associated with nucleus. Strikingly, when co-expressed in HFFs, GRA29 frequently formed smaller puncta from which SFP1 filaments radiated suggesting GRA29 can initiate SFP1 strand formation (Fig. 3C).

**Figure 3.**
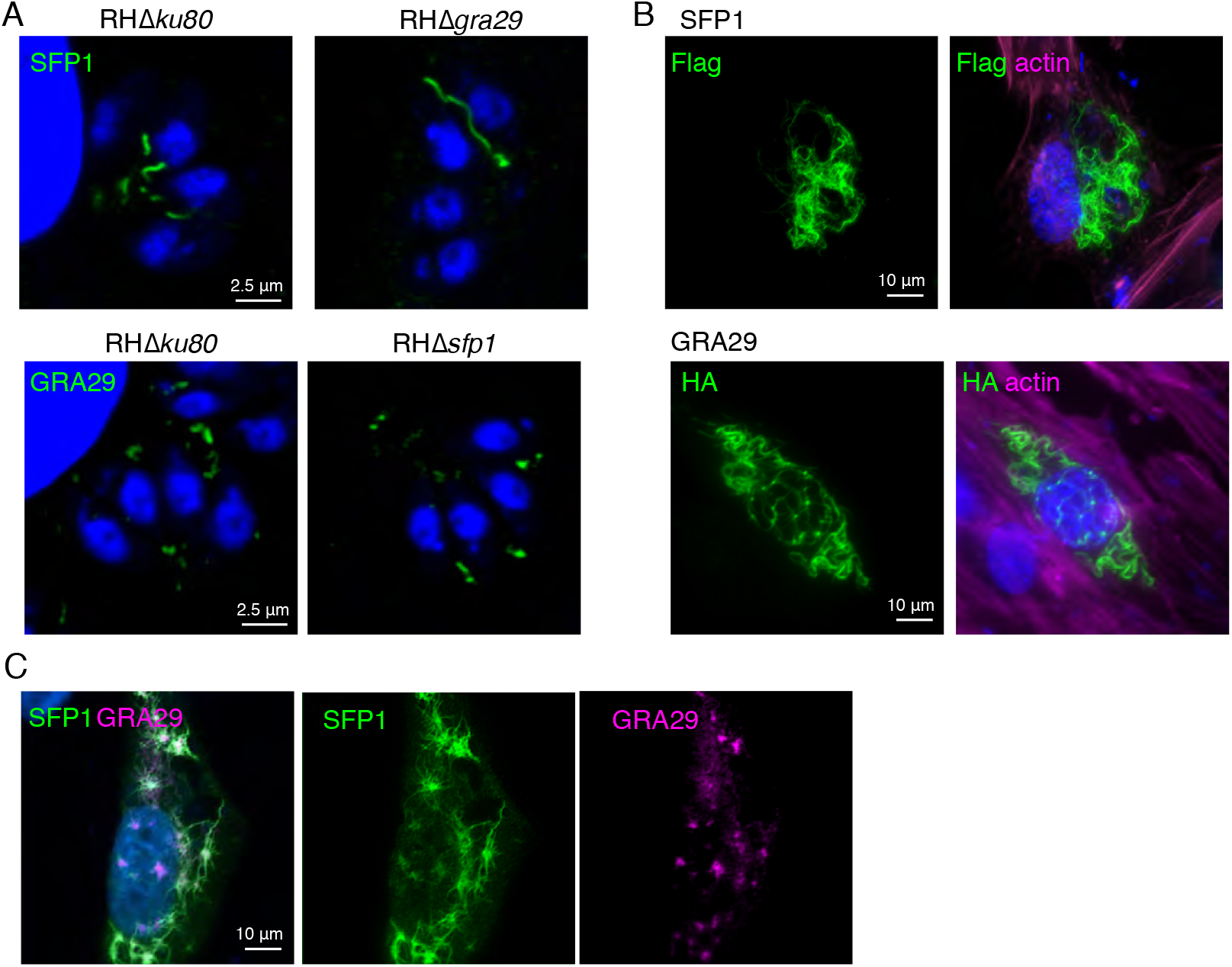
GRA29 regulates SFP1 strand formation. **a)** Fewer and longer SFP1 strands form in the absence of GRA29. Immunofluorescence analysis of HFFs infected with RHΔ*ku80*, RHΔ*sfp1* or RHΔ*gra29*. Anti-SFP1 (top panel), anti-GRA29 (lower panel) and DAPI (blue). **b)** SFP1 and GRA29 form strands when ectopically expressed. Immunofluorescence analysis of HFFs expressing N-terminally tagged SFP1 (Flag) or GRA29 (HA). Anti-Flag or HA antibodies (green), Phalloidin (magenta) to visualise F-actin, and DAPI (blue). **c)** HFFs co-expressing Flag-SFP1 and HA-GRA29 show SFP1 strands radiating from GRA29 foci. Immunofluorescence of co-transfected HFFs expressing Flag-SFP1 and HA-GRA29. Anti-SFP1 (green), anti-HA (magenta) and DAPI (blue).

These data indicate that both SFP1 and GRA29 can form filament-like structures in the absence of other *Toxoplasma* proteins. Although we were unable to confirm the interaction in co-immunoprecipitation or biochemical experiments (data not shown) the co-localisation of SFP1 and GRA29 even in this non-physiological context suggests that they can directly interact. Furthermore, the localisation pattern observed in HFFs and the extended SFP1 filaments in parasites lacking GRA29 suggest that GRA29 initiates SFP1 strand formation.

### The SFP1/GRA29 C terminal tail is required for strand formation

The change of localisation of GRA29 upon C-terminal HA-tagging and the cluster of phosphorylation sites in the C-termini of SFP1 and GRA29 suggested the C-terminal tail may play an important role in the regulation of strand-formation. To address this, we expressed truncated SFP1 and GRA29 lacking the unstructured C-terminal region in HFFs. In contrast to forming filaments under these conditions, both proteins appeared distributed throughout the HFF cytoplasm, with sparse foci of aggregation observed (Fig. 4A). This strongly suggested that the C-terminus of the protein plays an important role in filament formation. Consistently, during *Toxoplasma* infection with RHΔ*sfp1*+SFP1Δ*Ct* there was a lack of typical SFP1 strand formation in the PV (Fig. 4B). However, SFP1Δ*Ct* appeared aggregated within the PV, rather than dispersed, suggesting other PV resident proteins impact SFP1 distribution.

**Figure 4.**
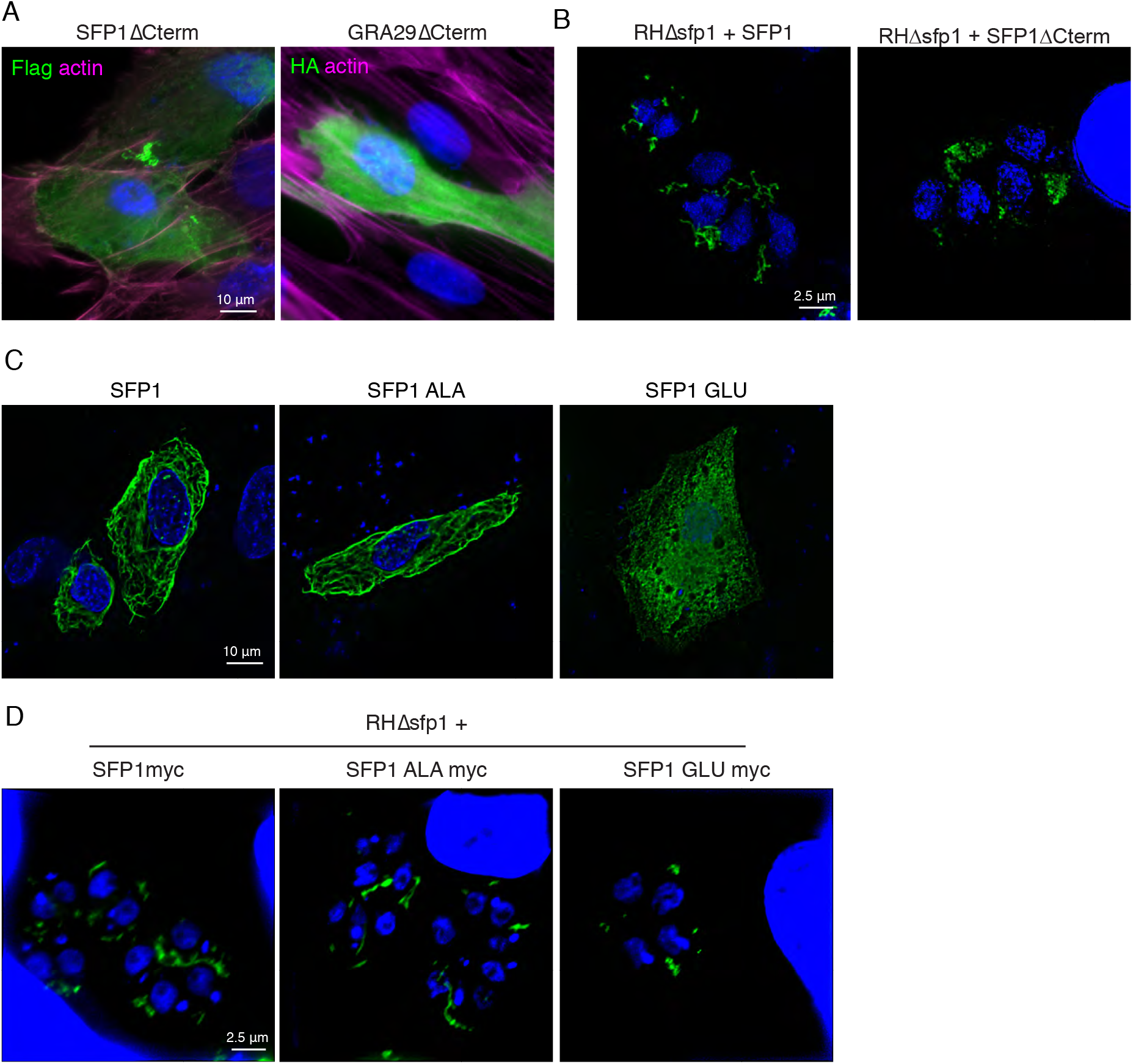
SFP1 and GRA29 C-termini regulate strand formation. **a)** Immunofluorescence analysis of HFFs expressing truncated Flag-SFP1 or HA-GRA29 lacking their disordered C termini. Anti-Flag or HA antibodies (green), Phalloidin (magenta) to visualise F-actin, and DAPI (blue). **b)** SIM analysis of HFFs infected with RHΔ*sfp1* expressing SFP1-myc or SFP1ΔCterm-myc. Anti-SFP1 antibodies (green) and DAPI (blue) **c)** SFP1 phosphomutants (preventing phosphorylation) does not disrupt strand formation. Immunofluorescence analysis of HFFs expressing Flag-SFP1, or Flag-SFP1 with the C-terminal phosphosites mutated to ALA or GLU. Anti-Flag antibodies (green) and DAPI (blue). **d)** Immunofluorescence analysis of HFFs infected with RHΔ*sfp1* expressing myc tagged SFP1, SFP1 ALA or SFP1 GLU. Anti-myc antibodies (green) and DAPI (blue).

Having established that the C-terminus of SFP1 is important for its subcellular organisation, we asked whether phosphorylation of SFP1 is important for filament formation. To investigate this, we first turned to expression of phosphomutants in HFFs, in which all phosphorylation sites in the SFP1 C-terminus were mutated to Alanine (SFP1_ALA) and phosphomimetics, in which all phosphorylation sites were mutated to Glutamic acid (SFP1_GLU). While SFP-ALA mutants displayed normal filaments in the HFFs, mimicking phosphorylation lead to an inability of SFP1 to form filaments. This indicates that phosphorylation may be a negative regulator of filament formation (Fig. 4C). This was replicated when SFP1_ALA and SFP1_GLU were expressed in RHΔ*sfp1* parasites (Fig. 4D), where extended strands were observed with phosphomutant but not phosphomimetic mutations.

If phosphorylation was preventing filament formation, we hypothesized that SFP1 and GRA29 expressed in HFFs should not be phosphorylated. To verify that assumption we immunoprecipitated SFP1 from HFFs and analysed the proteins by mass-spectrometry. Despite the presence of many non-modified peptides from SFP1, we observed no phosphopeptides (data not shown), indicating that indeed, SFP1 is not phosphorylated by human kinases when expressed in HFFs.

Collectively these data show that the C-termini of SFP1 and GRA29 are important for their regulation and that phosphorylation appears as a negative regulator with the non-phosphorylated protein forming strands.

### Secreted proteins are differentially phosphorylated in the chronic stage

To determine whether SFP1 and GRA29 are required *in vivo*, we generated gene KOs in the Type II strain Pru, and verified gene disruption by Western blot (Fig. S3A). Mice infected with PruΔ*sfp1* and PruΔ*gra29* parasites succumbed to infection similarly to WT PruΔ*ku80* parasites indicating neither protein is essential for parasite survival *in vivo* (Fig. S4B, C). During our experiments with the Pru strains, we observed that in some vacuoles SPF1 and GRA29 appeared distributed throughout the PV space instead of forming characteristic strands (Fig. 5A). Type II strains differ from the type I RH strain in that they more frequently convert to slow-growing bradyzoites in tissue culture. Bradyzoite cysts can be distinguished from tachyzoite vacuoles using fluorescently labelled lectin (DBA) that binds to the cyst wall (25). Parasites with a disperse SFP1/ GRA29 pattern also stained positive for the cyst-wall marker (Fig. 5A). Spontaneous conversion to bradyzoites occurs at low levels in standard tissue culture but high levels of conversion can be induced by pH stress (26). Under bradyzoite-inducing conditions parasite vacuoles were positive for the cyst wall marker CST1, and showed disperse SFP1 and GRA29 indicating that both proteins undergo a major change in localisation between the tachyzoite and the bradyzoite stages (Fig.5B).

**Figure 5.**
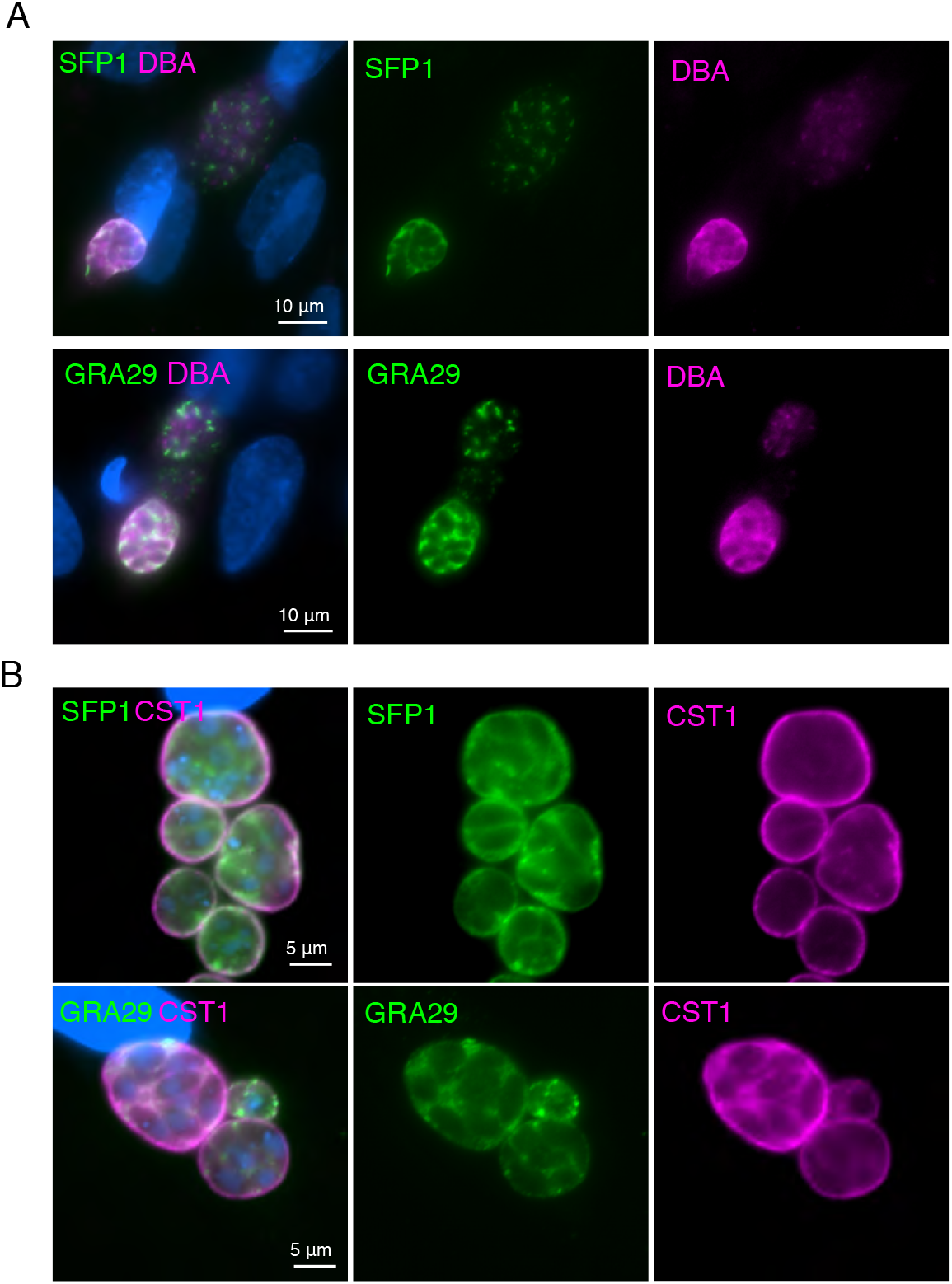
SFP1 and GRA29 disperse in chronic stage cysts. **a)** Immunofluorescence analysis of HFFs infected with PruΔ*hxgrpt* for 40h. Anti-SFP1 or anti-GRA29 (green), DBA (red) to stain the cyst wall and DAPI (blue). **b)** Immunofluorescence analysis of PruΔ*hxgrpt* infected HFFs grown under bradyzoite inducing conditions for 3 days. Anti-SFP1 or anti-GRA29 (green) with anti-CST1 (magenta) to visualise the cyst wall, and DAPI (blue).

We hypothesised that increased phosphorylation of SFP1 and GRA29 under bradyzoite conditions could lead to their dispersion. This phenomenon of redistribution in the bradyzoite cyst has been described for other GRAs and so we wanted to determine if these were also differentially phosphorylated. We therefore used comparative phosphoproteomics to determine differences in the phosphorylation of SFP1 and GRA29 and more broadly between acute (tachyzoite) and chronic (bradyzoite) conditions. Triplicate samples of *Toxoplasma* were grown in tachyzoite conditions for 27 h or bradyzoite conditions for 3 days (to give comparable numbers of parasites per vacuole) (Fig. 6A). We performed quantitative mass spectrometry to compare protein and phosphosite abundances between the two conditions using tandem mass tags (TMT) as described in (27, 28). First, we analysed differential protein abundance and identified 171 differentially regulated proteins with 103 proteins with higher abundance in bradyzoites and 68 with higher abundance in tachyzoites. As expected, known bradyzoite markers such as BAG1, ENO1, LDH2 and MAG1 were more abundant in the bradyzoite samples compared to the tachyzoite samples (Table S1 and Fig. S5).

**Figure 6.**
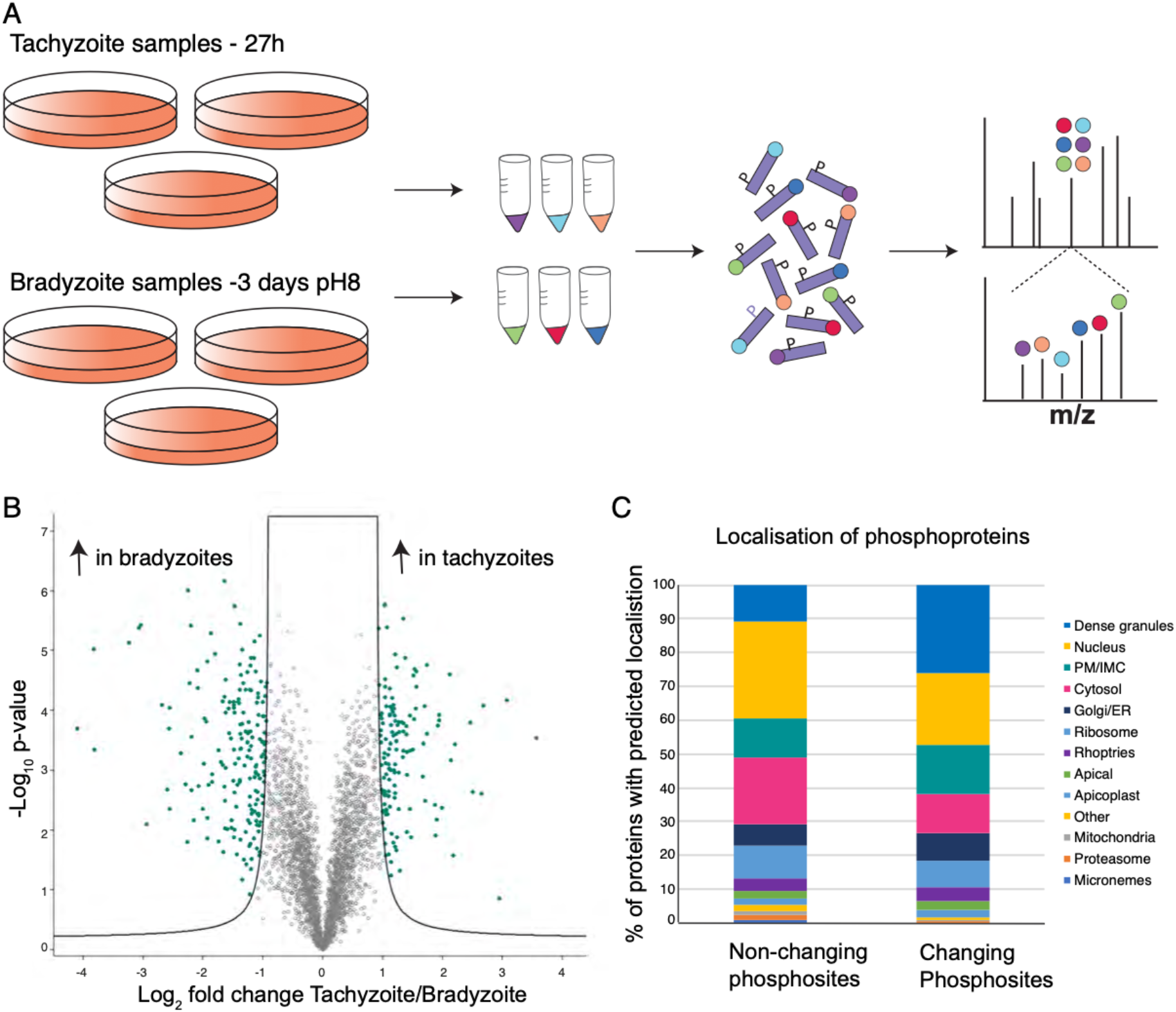
Secreted proteins are differentially phosphorylated in acute and chronic stages. **a)** Schematic of quantitative phosphoproteomics experiment. Peptides from triplicate tachyzoite and bradyzoites samples were labelled with tandem mass tags (TMT), enriched for phosphopeptides and analysed by mass spectrometry. **b)** Volcano plot showing differential phosphorylation between tachyzoite and bradyzoite conditions after normalisation to protein abundance. Each circle represents a phosphosite and those that change significantly (FDR 1%, −0.9> log_2_ fold change >0.9) are shown in green. **c)** Dense granule proteins are enriched in the subset of proteins that show significantly changing phosphorylation between tachyzoites and bradyzoites. Chart showing the predicted localisation of phosphoproteins using LOPIT (ToxoDB).

In the phosphopeptide enriched samples we quantified 7650 phosphorylation sites on 2235 *Toxoplasma* proteins. For 3730 of the phosphorylation sites we also obtained quantitative proteome data to which phosphorylation site changes were normalized. This allowed us to assess true differential phosphorylation rather than changes originating from differential protein abundance between conditions. After this normalisation step, we identified 337 phosphorylation sites that were significantly different between tachyzoite (144 sites more phosphorylated) and bradyzoite (193 sites more phosphorylated) stages (Fig. 6B, Table S1). The 337 phosphorylation sites are found on 170 proteins, 51 of which are predicted to be secreted (ROPs or GRAs in LOPIT dataset, ToxoDB). While GRAs made up 11% of proteins with non-changing phosphosites, they represented 26% of differentially phosphorylated proteins (Fig. 6C). This enrichment of GRAs in the subset of proteins that are differentially phosphorylated between tachyzoite and bradyzoite conditions indicates that these are indeed subject to extensive differential phosphorylation during stage conversion.

In this experiment we also identified phosphosites corresponding to the C-termini of SFP1 and GRA29, but they did not significantly change between conditions, (although a previously undetected N-terminal site on GRA29 was detected as more phosphorylated in bradyzoites). However, the GRAs that were detected as differentially phosphorylated included the IVN localised GRA2, GRA4, GRA6, GRA12, and PV membrane GRA1 and GRA5 (Table 1). We also detected differential phosphorylation of the cyst wall proteins CST1, CST3, CST4 and CST6. Furthermore, of the 38 recently identified putative cyst wall components (29), 24 were differentially phosphorylated suggesting phosphorylation could play an important role in the formation of this structure.

**Table 1.**
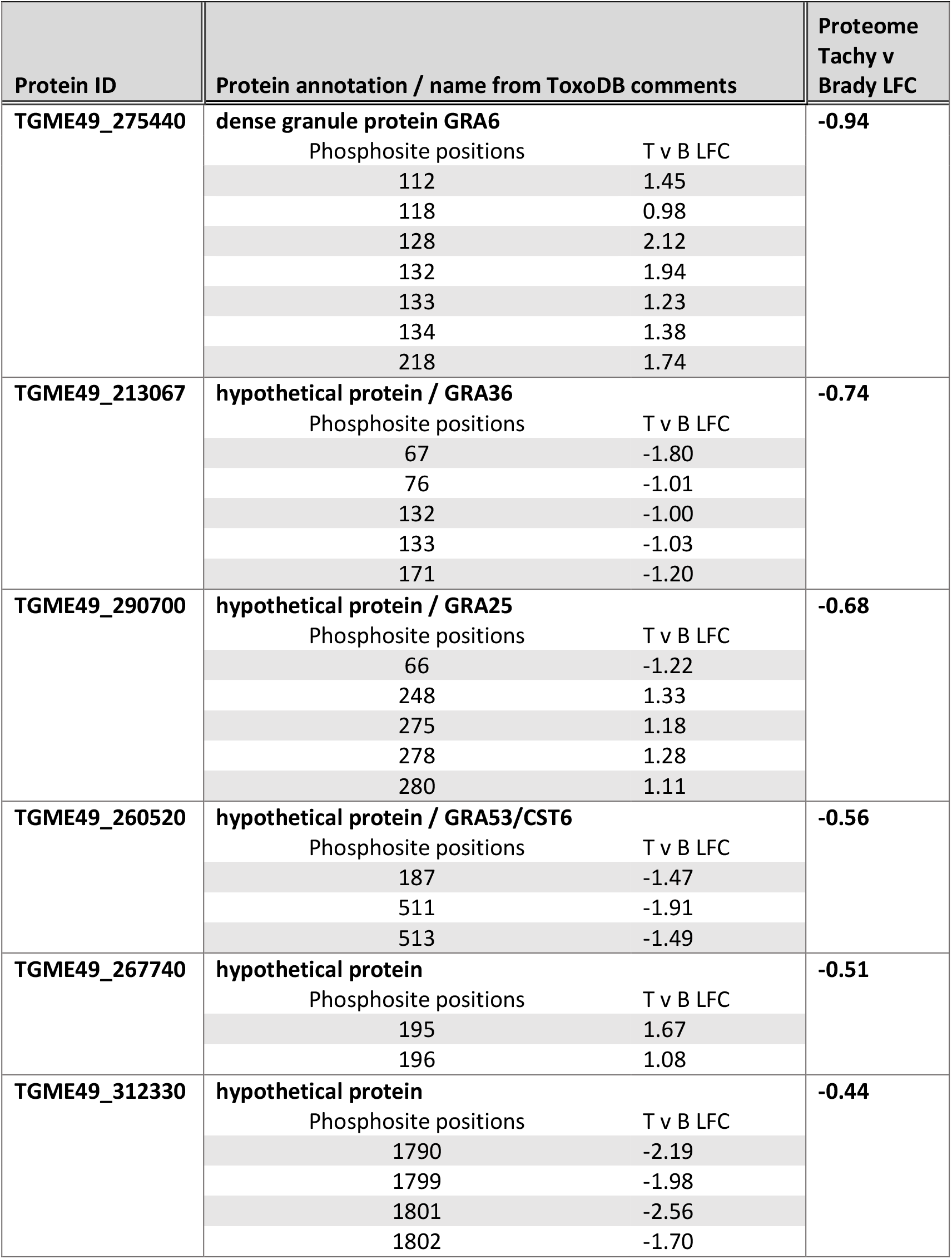
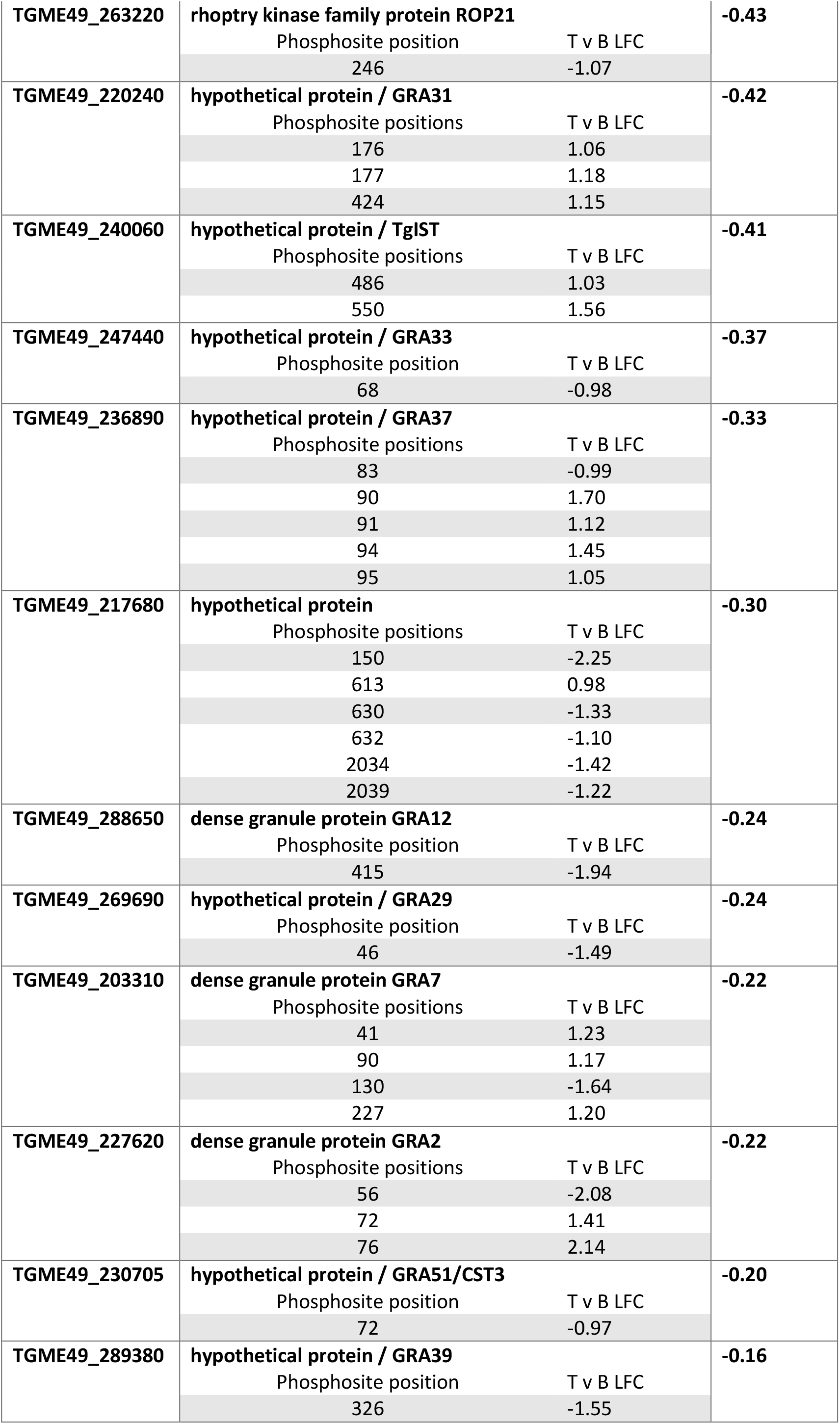
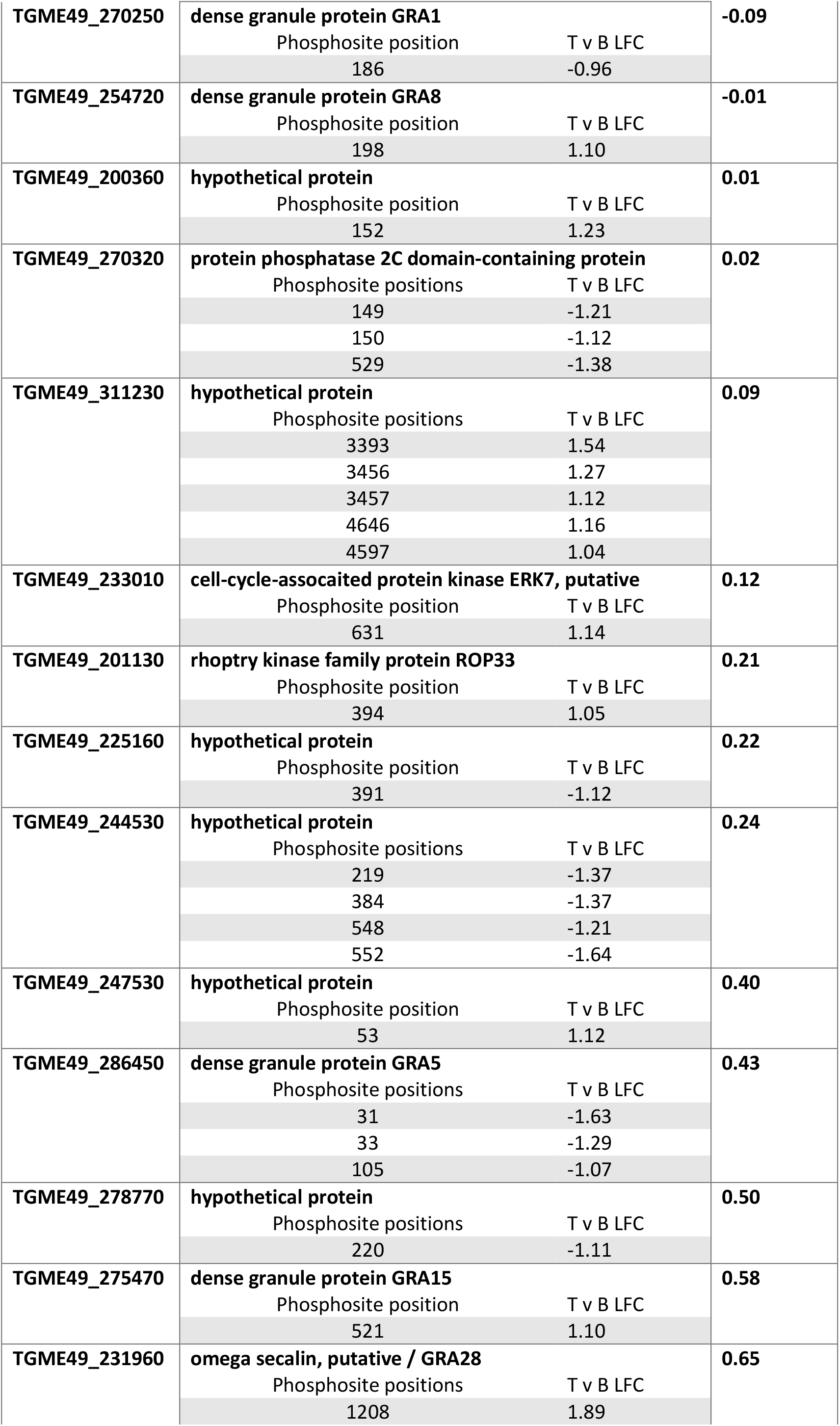
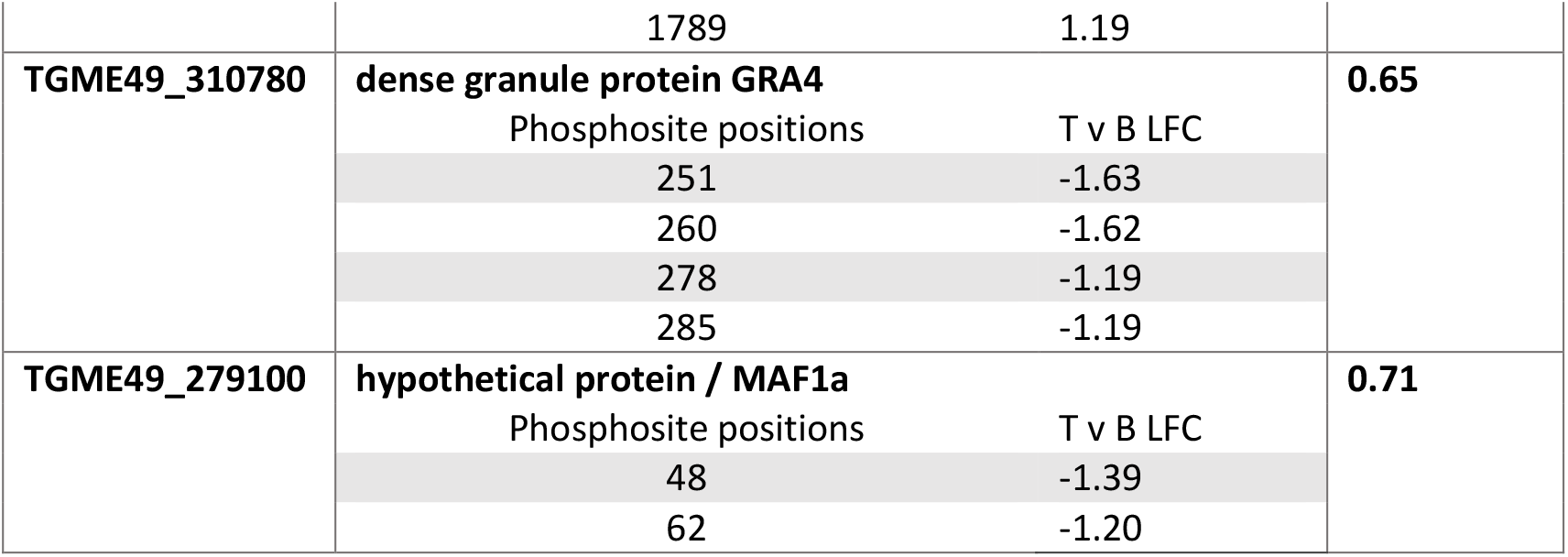
Differential phosphorylation of predicted dense granule proteins between tachyzoite and bradyzoites. Subset of Table S1 showing proteins predicted to localise to the dense granules with a Log_2_ fold change (LFC) in protein abundance <1. LFC for proteome and the normalised LFC for each phosphosite are shown. Negative scores indicate an enrichment in bradyzoites. Full details in Table S1.

| Protein ID | Protein annotation / name from ToxoDB comments | Proteome Tachy v Brady LFC |
| --- | --- | --- |
| TGME49_275440 | **dense granule protein GRA6** | **-0.94** |
|  | Phosphosite positions T v B LFC |  |
|  | 112 1.45 |  |
|  | 118 0.98 |  |
|  | 128 2.12 |  |
|  | 132 1.94 |  |
|  | 133 1.23 |  |
|  | 134 1.38 |  |
|  | 218 1.74 |  |
| TGME49_213067 | **hypothetical protein / GRA36** | **-0.74** |
|  | Phosphosite positions T v B LFC |  |
|  | 67 -1.80 |  |
|  | 76 -1.01 |  |
|  | 132 -1.00 |  |
|  | 133 -1.03 |  |
|  | 171 -1.20 |  |
| TGME49_290700 | **hypothetical protein / GRA25** | **-0.68** |
|  | Phosphosite positions T v B LFC |  |
|  | 66 -1.22 |  |
|  | 248 1.33 |  |
|  | 275 1.18 |  |
|  | 278 1.28 |  |
|  | 280 1.11 |  |
| TGME49_260520 | **hypothetical protein / GRA53/CST6** | **-0.56** |
|  | Phosphosite positions T v B LFC |  |
|  | 187 -1.47 |  |
|  | 511 -1.91 |  |
|  | 513 -1.49 |  |
| TGME49_267740 | **hypothetical protein** | **-0.51** |
|  | Phosphosite positions T v B LFC |  |
|  | 195 1.67 |  |
|  | 196 1.08 |  |
| TGME49_312330 | **hypothetical protein** | **-0.44** |
|  | Phosphosite positions T v B LFC |  |
|  | 1790 -2.19 |  |
|  | 1799 -1.98 |  |
|  | 1801 -2.56 |  |
|  | 1802 -1.70 |  |

|  |  |  |  |
| --- | --- | --- | --- |
| TGME49_263220 | **rophtry kinase family protein ROP21** |  | **-0.43** |
|  | Phosphosite position | T v B LFC |  |
|  | 246 | -1.07 |  |
| TGME49_220240 | **hypothetical protein / GRA31** |  | **-0.42** |
|  | Phosphosite positions | T v B LFC |  |
|  | 176 | 1.06 |  |
|  | 177 | 1.18 |  |
|  | 424 | 1.15 |  |
| TGME49_240060 | **hypothetical protein / TgIST** |  | **-0.41** |
|  | Phosphosite positions | T v B LFC |  |
|  | 486 | 1.03 |  |
|  | 550 | 1.56 |  |
| TGME49_247440 | **hypothetical protein / GRA33** |  | **-0.37** |
|  | Phosphosite position | T v B LFC |  |
|  | 68 | -0.98 |  |
| TGME49_236890 | **hypothetical protein / GRA37** |  | **-0.33** |
|  | Phosphosite positions | T v B LFC |  |
|  | 83 | -0.99 |  |
|  | 90 | 1.70 |  |
|  | 91 | 1.12 |  |
|  | 94 | 1.45 |  |
|  | 95 | 1.05 |  |
| TGME49_217680 | **hypothetical protein** |  | **-0.30** |
|  | Phosphosite positions | T v B LFC |  |
|  | 150 | -2.25 |  |
|  | 613 | 0.98 |  |
|  | 630 | -1.33 |  |
|  | 632 | -1.10 |  |
|  | 2034 | -1.42 |  |
|  | 2039 | -1.22 |  |
| TGME49_288650 | **dense granule protein GRA12** |  | **-0.24** |
|  | Phosphosite position | T v B LFC |  |
|  | 415 | -1.94 |  |
| TGME49_269690 | **hypothetical protein / GRA29** |  | **-0.24** |
|  | Phosphosite position | T v B LFC |  |
|  | 46 | -1.49 |  |
| TGME49_203310 | **dense granule protein GRA7** |  | **-0.22** |
|  | Phosphosite positions | T v B LFC |  |
|  | 41 | 1.23 |  |
|  | 90 | 1.17 |  |
|  | 130 | -1.64 |  |
|  | 227 | 1.20 |  |
| TGME49_227620 | **dense granule protein GRA2** |  | **-0.22** |
|  | Phosphosite positions | T v B LFC |  |
|  | 56 | -2.08 |  |
|  | 72 | 1.41 |  |
|  | 76 | 2.14 |  |
| TGME49_230705 | **hypothetical protein / GRA51/CST3** |  | **-0.20** |
|  | Phosphosite position | T v B LFC |  |
|  | 72 | -0.97 |  |
| TGME49_289380 | **hypothetical protein / GRA39** |  | **-0.16** |
|  | Phosphosite position | T v B LFC |  |
|  | 326 | -1.55 |  |
| TGME49_270250 | **dense granule protein GRA1** |  | **-0.09** |
|  | Phosphosite position | T v B LFC |  |
|  | 186 | -0.96 |  |
| TGME49_254720 | **dense granule protein GRA8** |  | **-0.01** |
|  | Phosphosite position | T v B LFC |  |
|  | 198 | 1.10 |  |
| TGME49_200360 | **hypothetical protein** |  | **0.01** |
|  | Phosphosite position | T v B LFC |  |
|  | 152 | 1.23 |  |
| TGME49_270320 | **protein phosphatase 2C domain-containing protein** |  | **0.02** |
|  | Phosphosite positions | T v B LFC |  |
|  | 149 | -1.21 |  |
|  | 150 | -1.12 |  |
|  | 529 | -1.38 |  |
| TGME49_311230 | **hypothetical protein** |  | **0.09** |
|  | Phosphosite positions | T v B LFC |  |
|  | 3393 | 1.54 |  |
|  | 3456 | 1.27 |  |
|  | 3457 | 1.12 |  |
|  | 4646 | 1.16 |  |
|  | 4597 | 1.04 |  |
| TGME49_233010 | **cell-cycle-associated protein kinase ERK7, putative** |  | **0.12** |
|  | Phosphosite position | T v B LFC |  |
|  | 631 | 1.14 |  |
| TGME49_201130 | **roptry kinase family protein ROP33** |  | **0.21** |
|  | Phosphosite position | T v B LFC |  |
|  | 394 | 1.05 |  |
| TGME49_225160 | **hypothetical protein** |  | **0.22** |
|  | Phosphosite position | T v B LFC |  |
|  | 391 | -1.12 |  |
| TGME49_244530 | **hypothetical protein** |  | **0.24** |
|  | Phosphosite positions | T v B LFC |  |
|  | 219 | -1.37 |  |
|  | 384 | -1.37 |  |
|  | 548 | -1.21 |  |
|  | 552 | -1.64 |  |
| TGME49_247530 | **hypothetical protein** |  | **0.40** |
|  | Phosphosite position | T v B LFC |  |
|  | 53 | 1.12 |  |
| TGME49_286450 | **dense granule protein GRA5** |  | **0.43** |
|  | Phosphosite positions | T v B LFC |  |
|  | 31 | -1.63 |  |
|  | 33 | -1.29 |  |
|  | 105 | -1.07 |  |
| TGME49_278770 | **hypothetical protein** |  | **0.50** |
|  | Phosphosite positions | T v B LFC |  |
|  | 220 | -1.11 |  |
| TGME49_275470 | **dense granule protein GRA15** |  | **0.58** |
|  | Phosphosite position | T v B LFC |  |
|  | 521 | 1.10 |  |
| TGME49_231960 | **omega secalin, putative / GRA28** |  | **0.65** |
|  | Phosphosite positions | T v B LFC |  |
|  | 1208 | 1.89 |  |
|  | 1789 | 1.19 |  |
| TGME49_310780 | **dense granule protein GRA4** |  | **0.65** |
|  | Phosphosite positions | T v B LFC |  |
|  | 251 | -1.63 |  |
|  | 260 | -1.62 |  |
|  | 278 | -1.19 |  |
|  | 285 | -1.19 |  |
| TGME49_279100 | **hypothetical protein / MAF1a** |  | **0.71** |
|  | Phosphosite positions | T v B LFC |  |
|  | 48 | -1.39 |  |
|  | 62 | -1.20 |  |

It is worth noting that we also observed differential phosphorylation on proteins important for DNA regulation. This includes the AP2 domain transcription factors AP2VIII-2 and AP2XII-4, and 32 predicted nuclear proteins. These may be important for transcriptional regulation during stage conversion, but their role is not further discussed here.

Collectively these data show that SFP1 and GRA29 undergo a dramatic change in localisation between tachyzoite and bradyzoite conversion but that their phosphorylation is transient or not required for this transition, at least at the experimental timepoint selected herein. Furthermore, we show widespread changes in phosphorylation state in a substantial number of secreted proteins, indicating the activity of stage-specific kinases and phosphatases in regulating PV resident proteins.

## DISCUSSION

Here we have analysed the localisation of two secreted phosphoproteins, SFP1 and GRA29 that re-localise during stage conversion to bradyzoites. Both proteins form stranded structures in the tachyzoite PV, that are distinct from the IVN and actin filaments previously observed. We did not identify a function of SFP1 and GRA29 during the lytic cycle or *in vivo* infections, preventing us from assessing the impact of their phosphorylation in *Toxoplasma* biology. It could be that the proteins have a subtle effect on growth that we have not been able to measure, or that they are important in another host-species, parasite stage or genetic background we have not tested here. However, neither SFP1 nor GRA29 showed a reduction in fitness *in vivo* (30) in our recent more sensitive *in vivo* CRISPR screen, supporting a non-essential role under these conditions.

We show the importance of the C-termini of both proteins for strand formation. Using mutational analysis, we show that SFP1 phosphorylation is not required for filament formation and that mimicking phosphorylation disrupts filament formation. As such, mimicking phosphorylation mirrored the phenotype observed with removal of the C-terminal tail. This suggests that SFP1 and GRA29 form oligomers dependent on their un-phosphorylated C-termini. Phosphorylation negatively regulates the intrinsic behaviour of SFP1 and GRA29 and abrogates their ability to form multimers. However, as both proteins where identified to be phosphorylated after secretion into the PV (19) where, as we show here, they form fibril-like structures, we assume that their phosphorylation is a dynamic process, potentially leading to the regulation of their length and location. It could also be that phosphorylated and non-phosphorylated SFP1 and GRA29 localize to different structures in the PV. How both proteins re-localise during stage conversion is unclear. While mimicking phosphorylation appears sufficient to promote dispersal of SFP1 and GRA29 in HFFs, this is not observed when expressed in tachyzoites, where an aggregation is observed. This indicates that other proteins, or post-translational modifications are involved in SFP1/ GRA29 regulation in parasites upon stage conversion.

Recently, the PV localised kinase WNG1/ROP35 was shown to phosphorylate several GRAs, including GRA6, and to regulate their membrane association (20). As many GRAs form complexes during trafficking (31), WNG1 phosphorylation is hypothesised to release them from association with a chaperone, freeing them to associate with membranes. As such, WNG1 dependent phosphorylation of GRAs was shown to promote their normal behaviour and localisation. We now propose that in addition to WNG1 phosphorylation promoting release and membrane association of GRAs, phosphorylation of secreted proteins can negatively regulate their behaviour. In this way, the parasite could use phosphorylation by kinases within the PV lumen to regulate PV resident proteins. This would allow the parasite to tailor its niche and respond to differing conditions or requirements.

Several secreted kinases localise within the PV lumen and are upregulated during the chronic stages of infection (for example ROP21, ROP27, ROP28, and indeed WNG1/ROP35). Importantly, secreted kinases have been shown to contribute to cyst formation in mice indicating that phosphorylation within the PV/forming cyst is required for cyst development. ROP21/27/28 contribute to the parasite establishing chronic infection but their function and targets remain unknown (32). Additionally, a triple knockout of ROP38/29/19 showed a severe defect in cyst formation (33). Here we expand our knowledge of cyst formation by reporting the differential phosphorylation of proteins within the forming cyst. Our data also identifies phosphorylation that is reduced upon stage conversion. The expression of many ROP family kinases is reduced in the chronic stage of infection, which could account for some changes, but it is likely that secreted phosphatases are contributing to the differential phosphorylation. The necessity of deleting several ROPs to observe strong phenotypic effects in bradyzoites may reflect that the system is flexible with a high degree of redundancy. Alternatively, the combined effect of multiple kinases could allow the parasite to integrate several signals leading to stage differentiation.

PV resident GRAs re-localise as the parasite converts to the chronic stage and remodels the PV to form the cyst. Guevera *et al.*, recently described the dynamic nature of this re-localisation over the course of cyst formation (for GRA1, 2, 4, 6, 9, 12) (21). Furthermore, they showed that IVN localised GRAs were required for normal formation of the cyst wall. Here we show that these GRAs, with the exception of GRA9 (which is upregulated in bradyzoites) are differentially phosphorylated in the acute and chronic stages suggesting phosphorylation could play a role in their regulation. Interestingly, GRA2 showed phosphorylation sites that were enriched in each condition; two sites within a predicted amphipathic helix were more phosphorylated in tachyzoite conditions, while a third site was more phosphorylated in the bradyzoite samples. It is tempting to speculate that dual phosphorylation within the polar side of the amphipathic helix (a1) of GRA2 could alter its membrane binding. Further work assessing phosphorylation of secreted proteins over time is required to decipher the impact of this modification and the role of both kinases and phosphatases in remodelling the PV. However, the regulation of proteins after parasite secretion adds an intriguing extra layer of complexity to pathogen effector biology and establishing *Toxoplasma* chronic infection.

## MATERIALS AND METHODS

### Cell culture and parasite strains

Primary human foreskin fibroblasts (HFFs) (ATCC) were maintained in Dulbecco’s modified Eagle’s Medium (DMEM) with 4.5g/L glucose and GlutaMAX-1 (Gibco) supplemented with 10% foetal bovine serum (FBS) at 37°C and 5% CO_2_. T. *gondii* RHΔ*ku80*::diCre (34), RHΔ*gra2* (35), PruΔ*ku80 (36)* and PruΔ*hxgprt* (gift from Dominique Soldati, as in (37)) parasites were maintained by growth in confluent HFFs and passaged every 2-3 days. RH strains were seeded 24 h, and Pru strains 48 h, before transfections or infection experiments.

### Sequence analysis

Gene sequences and predicted protein localisations (LOPIT dataset) were obtained from ToxoDB. Alignments were generated using ClustalW and schematics using DOG2.0 (38). Protein sequences were analysed by Multicoil2, (39)) and Globplot, (40) to identify coiled coil and unstructured regions respectively. Protein schematics were generated

### Plasmid construction

All primers are shown in Supplementary File 1. Vectors were generated by inverse PCR using KOD Hot-start polymerase (Novagen) or by Gibson assembly in a homemade reaction master mix (100 mM Tris-Cl pH 7.5, 50 mg/mL PEG-8000, 10 mM MgCl_2_, 10 mM dithiothreitol, 0.2 mM each of four dNTPs, 1 mM NAD with 0.8U T5 exonuclease (NEB), 0.5 U Phu DNA polymerase (NEB) and 80 U Taq DNA ligase (NEB)). Gibson reactions were incubated for 60 min at 50°C. All vectors were verified by sequencing. *sfp1* and *gra29* sequences were amplified from *Toxoplasma* gDNA prepared from 1E7 RHΔ*ku80*::diCre parasites using Qiagen Blood and Tissue DNA extraction kit, unless otherwise stated.

To generate pSFP1, the TGGT1_289540 promoter and coding sequence were amplified using primers 1 & 2 and inserted in to HindIII/PacI digested pGRA (12). A C terminal myc tag was inserted by inverse PCR using primers 3 and 4. pSFP1ΔCterm was generated by inverse PCR on pSFP1 with primers 4 & 5. To generate phosphomutant versions of pSFP1, initially a silent ApaI site was inserted into pSFP1 (at nt2529) by inverse PCR with primers 6 & 7. 400bp synthetic DNA fragments (IDT, Supplementary file 1) encoding SFP1-myc C-terminal or versions with Ser/Thr to ALA, or Ser/Thr to GLU mutations were mixed with ApaI/PacI digested pSFP1 ApaI in Gibson reaction mix. pGRA29-HA was generated by amplifying GRA29 promoter and coding sequence using primers 8 & 9 and inserting the amplicon in to HindIII/NcoI digested pGRA.

For pcDNA-NTAP-Flag SFP1 constructs, the SFP1 coding sequence excluding the signal peptide was amplified with primers 10 & 11 for full length or primers 10 & 12 for C terminal truncated versions. For phosphosite mutant versions primers 10 & 11 were used with pSFP1 ALA or pSFP1 GLU as the template. All amplicons were inserted into NotI/EcoRI digested pcDNA-NTAP, a pcDNA3.1 derivative for expression with 3x Flag tag, TEV cleavage site and calmodulin binding peptide. For pcDNA-HA-GRA29 construct, GRA29 was initially amplified with primers 13 & 14 and inserted into NotI/EcoRI digested pcDNA-NTAP. The Flag tag was removed and an N-terminal HA tag inserted by inverse PCR with primers 15 & 16. pcDNA-GRA29ΔCterm was generated by inverse PCR with primers 17 & 18.

gRNA sequences targeting the 5’ and 3’ of *sfp1/gra29* were inserted into pSAG1::CAS9-U6::gUPRT (41) by inverse PCR with primers 19 & 20-23. Dual gRNA pSAG1::CAS9 vectors were generated by amplifying 3’ gRNA expression locus with primers 24 & 25 and inserting the region into XhoI/PacI digested pSAG1::CAS9 carrying the 5’ gRNA.

To generate the SFP1 bacterial expression plasmid, pET-28a-SFP1, SFP1 lacking its amino terminal signal peptide was amplified using primers 26 & 27. The amplicon was cloned into *BamHI* and *NdeI* digested pET-28a(+) plasmid (Merck).

### Parasite transfection

Parasites were released from HFFs by syringe lysis, pelleted at 650 × g and resuspended in cytomix (10mM KPO4, 120mM KCl, 0.15mM CaCl2, 5mM MgCl_2_, 25mM Hepes, 2mM EDTA). ^~^5E6 parasites were combined with 40 ug DNA in 400ul cytomix in a 2mm cuvette and electroporated using a Bio-Rad Gene Pulser Xcell at 1500 V, and 25 uF. Repair templates were linearised by overnight digestion, purified by phenol-chloroform precipitation and resuspended in H2O. 24h post transfection 50 μg/ml Mycophenolic acid (Merck) and Xanthine (Sigma) (M/X) were added to select for transfectants.

### Generating parasite lines

RHΔ*ku80* lacking SFP1 was generated by transfecting parasites with pSAG1-CAS9sgSFP1 targeting the 5’ end of the gene (starting 84nt after ATG) with a 183nt repair template to remove the PAM site and introduce STOP codons (seq in supp file?). Parasites expressing the transfected Cas9-GFP were sorted by flow cytometry into 96 well plates containing HFF monolayers using a BD Influx cell sorter (BD Biosciences). This line was used for complementing pSFP1myc constructs. Alternatively, *gra29* and *sfp1* were disrupted in RHΔ*ku80* and PruΔ*ku80* backgrounds by co-transfecting a double gRNA pCAS9 vector targeting the 5’ and 3’ of the gene and a synthetic repair construct with 500 nt homology arms and HXGPRT-T2A-mCherry cassette (GeneArt, Life Technologies). Transfectants were selected by the addition of M/X, and clones were screened by immunofluorescence and verified by western blot.

### Antibody generation

N-terminal 6xHis tagged SFP1 recombinant protein was expressed form pET-28a-SFP1 in *Escherichia coli* BL21 cells. His-SFP1 was purified using Ni-NTA affinity purification under native conditions using the standard manufacturer’s protocol (Qiagen). Recombinant His-SFP1 was used to immunize female New Zealand white rabbits (Covalab) for generation of polyclonal antibodies. To generate the mouse anti-SFP1 antibody, phosphopeptides corresponding to the C-terminal of SFP1 were conjugated to Keyhole limpet hemocyanin and used to immunise 4 mice (Covalab). Day 53 sera showed reactivity against SFP1. While mice were immunized with phosphorylated isoforms, the antibodies recognise non-phosphorylated peptides.

SFP1 peptides used for immunisation. Residues in bold were phosphorylated.

*NH2-C-GSVA**S**GFRG -CONH2*

*NH2-C-GFRG**S**MA**S**GLFP – CONH2*

*NH2-C-LRGA**S**VAG**S**LGG -CONH2*

*NH2-C-GGVG**S**RLGG-CONH2*

*NH2-C-FAGA**S**MGRG-CONH2*

### Immunoprecipitation

HFFs were infected with RHΔ*ku80* or RHΔ*ku80* +SFP1::myc in a T25 flask for 24h before lysis in 1ml RIPA buffer (Thermoscientific) supplemented with protease (complete mini, Roche) inhibitors. Samples were clarified by centrifugation and incubated with anti-Myc tag antibody conjugated agarose (Millipore, Cat. no. 16-219) for 3h. The non-bound fraction was removed and the proteins precipitated with acetone at −20 °C. Agarose bead complexes were washed in RIPA buffer followed by Tris buffer saline and proteins eluted by boiling in Laemmli buffer (60 mM Tris-HCl pH6.8, 1% SDS, 5% glycerol, 5% b-mercaptoethanol, 0.01% bromophenol blue).

### Western blotting

*Toxoplasma* lysates or IP samples were separated by SDS-PAGE, transferred onto nitrocellulose (BioRad) and probed with appropriate antibodies in PBS with 3% milk and 0.1% Tween20 (Sigma-Aldrich). Rabbit and mouse anti-SFP1 serum were used at a dilution of 1:1000, rabbit anti-GRA29 serum at 1:1000 (ref CRISPR paper?), mouse anti-GRA1 at 1: 1000 (gift from Jean-Francois Dubremetz), mouse anti-GRA2 at 1:1000 (Biotem BIO.018.5), mouse anti-Myc at 1:1000 (Millipore, 05-724), and rat anti-HA at 1:1000 (Roche, 11867423001). After washing, the blots were incubated with HRP-conjugated secondary antibodies; anti-rabbit at 1:20,000 (Insight Biotechnology, Cat no. 474-1506), anti-mouse at 1:3,000 (Insight Biotechnology, Cat no. 474-1506) or anti-rat at 1:4000 (Life technologies, Cat. no 629520). The blots were visualised using ECL Western Blotting Detection Reagent (GE Healthcare) for chemiluminescence imaging on an Amersham Imager 600 (GE-Healthcare).

### Parasite infections

For widefield, confocal or EM imaging HFFs were seeded on 13mm #1.5 coverslips in 24 well plates. Confluent monolayers were infected at a multiplicity of infection (MOI) of 1 for 16-24 h. For Cytochalisin D treatment infected cells were incubated with 1 μM Cytochalasin D or DMSO for 1h before fixation. Samples were washed with PBS before fixation in 3% formaldehyde (FA) for 15 min. For EM analysis, infected cells were fixed in 4% FA for 15min followed by 2.5% glutaraldehyde/4% FA for 30min and storage in 1% FA.

### Conversion to bradyzoites

To induce parasite switching to bradyzoites, HFF monolayers were infected at an MOI of 0.5 for 3.5h then washed four times with PBS before adding switch media (RPMI (Sigma, R4130) with 1% FBS, pH 8.1) and incubating at 37°C and ambient CO_2_ for 3 days. Cells were fixed in 3% FA, quenched in 50mM ammonium chloride and permeabilised in 0.2% triton, 0.1% glycine 0.2% BSA in PBS for 20 mins on ice.

### Intracellular fluorescent protein ingestion assay

Intracellular fluorescent protein ingestion was assessed as described in McGovern et al. (42). Briefly, 70-80% confluent CHO-K1 cells were transfected in 35 mm dishes using X-TREMEGENE 9 Transfection Reagent (Roche, 6365787001). Each dish was transfected with 2 μg of pCDNA 3.1 Venus plasmid using the using a 3:1 ratio of plasmid to transfection reagent in Opti-MEM (Gibco, 31985062) and a total final volume of 200 μl. The CHO K1 cells were then incubated overnight at 37°C and synchronously infected with *T. gondii* parasites using the ENDO Buffer Method at 18 to 24h post-transfection (43). *T. gondii* parasites were treated with 1 μM LHVS or equal volume of DMSO for 48h prior to synchronous invasion, replacing the LHVS and DMSO-treated media every 6-18h, and also during the 3h incubation period after synchronous invasion. *T. gondii* parasites were purified at 3h post-invasion as previously described (16), pelleted by centrifugation at 1000xg for 10 min at 4°C, and subjected to a protease protection assay by resuspension in 250 μl of freshly prepared 1 mg/mL Pronase (Roche, 10165921001)/0.01% Saponin/PBS at 12°C for 1h. The parasites were then pelleted by centrifugation at 1000*xg* for 10 min and washed 3 times in ice cold PBS before depositing on Cell-Tak (Corning, 354240) coated chamber slides. Parasites were fixed with 4% paraformaldehyde imaged with an AxioCAM MRm camera-equipped Zeiss Axiovert Observer Z1 inverted fluorescence microscope. Ingestion of host cytosolic Venus was scored manually as Venus-positive or Venus-negative without sample blinding.

### HFF transfection

HFFs were seeded at a density of 7.5 x10^4^ cells per well on a 13mm glass coverslip 24h before transfection with Lipofectamine 2000 (Thermofisher). For each well 1.5 μl Lipofectamine 200 was mixed with 800 ng plasmid DNA in 100 μl Optimem (Thermofisher). Cells were fixed in 3% FA 18h post transfection.

### Immunofluorescence analysis

For immunofluorescence analysis cells were quenched in 50mM ammonium chloride for 15min. Transfection experiments and initial RHΔ*ku80* +SFP1::myc or RHΔ*ku80* +GRA29::HA infections were permeabilised for 3min in 0.1% triton and blocked with 2% BSA in PBS. For subsequent infection experiments samples were permeabilised and probed in 0.05 % saponin, 0.2% fish gelatin in PBS. Infected or transfected cells were probed with mouse anti-Myc (1:1000, Millipore 05-724), rat anti-HA (1:1000, Roche, 11867423001), mouse anti-Flag (1:1000, Sigma F1804), rabbit anti-GRA29 (1:4000,(30)), rabbit anti-SFP1 serum (1:1000), mouse anti-SFP1 (1:200), or mouse anti-CST1 (1:100, gift from Louis Weiss) followed by Alexafluor488 or 564 conjugated secondary antibodies (1:2000, Life Technologies). Additionally, cells were labelled with DAPI (1:1000 of 5mg/ml, Sigma Aldrich), Alexa647 conjugated Phalloidin (1:100, Invitrogen A22287) or Rhodamine conjugated Dolichos Biflorus Agglutinin (1:200, Vector Labs RL-1302). Antibodies/stains were diluted in 2% BSA PBS and samples were incubated for 1h and coverslips were subsequently washed and mounted with ProLong Gold anti-Fade moutant (Thermofisher, P36934). Localisations were visualised on a Ti-E Nikon microscope or Leica SP5 Confocal.

### Super resolution microscopy

For super resolution microscopy 24 × 24 #1.5H coverslips were washed in 1M HCl overnight, rinsed in water before coating with 1% (w/v) porcine gelatin (Sigma, G1890) in 6 well plates. Confluent HFF monolayers were infected with an MOI of 0.5 and centrifuged at 180 × g for 3 min to synchronise infection. Samples were fixed and processed as above except that antibody dilutions were passed through 0.45 μM filters and an additional final fixation step (10 min, 2 % FA) was included after secondary antibody incubation. Super-resolution microscopy imaging was performed using an OMX v3 structured illumination microscope (Applied Precision/GE Healthcare) equipped with 405, 488 and 592.5 nm diode lasers, electron multiplying CCD cameras (Cascade II 512 × 512, Photometrics) and a 100×/1.40 NA PlanApochromat oil-immersion objective lens (Olympus). 3D-SIM image stacks were reconstructed and aligned using softWoRx (Applied Precision/GE Healthcare).

### Correlative Light and Electron Microscopy

HFFs were seeded on a 3.5 cm dish with gridded coverslip (MatTek) and infected, fixed and processed as described above. Regions of interest with GRA29-HA cage structures were selected and imaged by SIM. Tiled images of the surrounding area were generated by Confocal imaging to aid in identifying the region by EM. The cells were then fixed and prepared for EM as described below. The light and electron microscopy overlay was generated using the TurboReg plugin of Fiji software, with an affine transformation using the parasite nuclei and apicoplasts as landmarks, and by selecting high intensity pixels in Photoshop using the magic wand tool with tolerance set to 40.

### EM sample preparation and imaging

Cells were fixed in 2.5% glutaraldehyde, 4% formaldehyde in 0.1 M phosphate buffer (PB) at room temperature for 30 min, post-fixed in 1% reduced osmium tetroxide for 1 h, stained with 1% tannic acid in 0.05 M PB for 45 min, and quenched with 1% sodium sulphate in 0.05 M PB for 5 min. For CLEM experiments, the gridded coverslip was then removed from the MatTek dish. After washing in PB and water, the cells were dehydrated in 70, 90, and 100% ethanol (2×5 min each) before infiltration with a 1:1 mix of propylene oxide and Epon resin (Taab Embed 812) for 1 h and then 2×90 min with Epon resin. Finally, coverslips were embedded in resin for 24 h at 60ºC, sectioned (70–75 nm), and stained with lead citrate. Images were acquired using a Tecnai Spirit Biotwin (Thermofisher Scientific) transmission electron microscope.

### *In vivo* infections

C57BL/6 (wild type) mice were bred and housed under pathogen-free conditions in the biological research facility at the Francis Crick Institute in accordance with the Home Office UK Animals (Scientific Procedures) Act 1986. All work was approved by the UK Home Office (project license PDE274B7D), the Francis Crick Institute Ethical Review Panel, and conforms to European Union directive 2010/63/EU. All mice used in this study were male and between 7- to 9-week-old.

Mice were infected with PruΔ*ku80*, PruΔ*sfp1* or PruΔ*gra29* tachyzoites by intraperitoneal injection (i.p.) with either 3E5 or 5E4 parasites in 200 μl PBS on day 0 to assess mouse survival. Mice were monitored and weighed regularly for the duration of the experiments.

## Mass spectrometry

### SFP1 transfection and Immunoprecipitation

2.8×10^6^ HFFs were seeded in a 15 cm dish 24 h before transfection with 44 ug pcDNA-SFP1 using Lipofectamine 3000 (Thermofisher) as per manufacturer’s instructions. 18h post transfection, cells were lysed on ice with 2 ml Immunoprecipitation (IP) buffer (Thermoscientific) for 30min, passed through a 23 G needle and clarified by centrifugation. The supernatant was incubated with 60ul of polyclonal SFP1 antibody coated Fast flow protein G sepharose (GE Healthcare). Sepharose bead complexes were washed in IP buffer followed by Tris buffer saline and proteins eluted by boiling in Laemmli buffer.

### In-gel digestion and LC-MS/MS

Eluted proteins were separated by sodium dodecyl sulfate polyacrylamide gel electrophoresis (SDS-PAGE), until the running front had migrated approximately 2 cm into the gel (12% NuPAGE, Invitrogen), and stained with colloidal Coomassie (InstantBlue, Expedeon). After excision of 8 horizontal gel slices per lane, proteins were in-gel digested with trypsin (Promega/Pierce) using a Janus liquid handling system (Perkin Elmer). Tryptic peptides were analyzed by liquid chromatography-mass spectrometry (LC–MS) using an Orbitrap Velos mass spectrometer coupled to an Ultimate 3000 uHPLC equipped with an EASY-Spray nanosource (Thermo Fisher Scientific) and acquired in data-dependent mode.

### MS Data Processing and Analysis

The data were searched against *Toxoplasma gondii* and *Homo sapiens* (both Uniprot) databases using the Andromeda search engine. Raw data were processed with MaxQuant (version 1.5.2.8). Cysteine carbamidomethylation was selected as a fixed modification. Methionine oxidation, acetylation of protein N-terminus and phosphorylation (S, T, Y) were selected as variable modifications. The enzyme specificity was set to trypsin with a maximum of 2 missed cleavages. The datasets were filtered on posterior error probability to achieve a 1% false discovery rate on protein, peptide and site level. Data were further analysed with Perseus (version 1.5.0.9). The data were filtered to remove common contaminants and IDs originating from reverse decoy sequences and only identified by site. The log10 values for peptide intensities were then determined and all SFP1 peptides (58% sequence coverage) analysed based on their modification state. No phosphorylated peptides were found.

## Tachyzoite v bradyzoite phosphoproteome

### Infection conditions

Confluent HFF monolayers were grown in 15 cm dishes and infected with PruΔ*hxgprt*. For tachyzoite samples three dishes were infected with 1.3×10^7^ parasites and incubated for 27h. For bradyzoite samples three plates were infected 1.1 x10^7^ parasites for 3.5 h before extensive washing and the addition of switch media (described above). Bradyzoite infections were incubated as above for 3 days with fresh switch media added daily.

### Cell lysis and protein digestion

Lysis was performed in ice cold 8 M urea, 75 mM NaCl, 50 mM Tris, pH 8.2, supplemented with protease (complete mini, Roche) and phosphatase (Phos Stop, Roche) inhibitors. Lysis was followed by sonication to reduce sample viscosity (30% duty cycle, 3 × 30 seconds bursts, on ice). Protein concentration was measured using a BCA protein assay kit (Pierce). Lysates (1mg each) were subsequently reduced with 5 mM DTT for 30 min at 56 °C and alkylated in the dark with 14 mM iodoacetamide for 30 min at RT. Following iodoacetamide quenching with 5 mM DTT for 15 min in the dark, lysates were diluted with 50 mM ammonium bicarbonate to < 4M urea, and digested with LysC (Promega) for 2-3h at 37 °C. Lysates were further diluted with 50 mM ammonium bicarbonate to <2M urea and digested with trypsin (Promega) overnight at 37°C. After digestion, samples were acidified with trifluoroacetic acid (TFA) (Thermo Fisher Scientific) to a final concentration of 1% (v/v). All insoluble material was removed by centrifugation and the supernatant was desalted on Sep-Pak C18 cartridges (Waters).

### TMT labelling

Samples were dissolved at 1 mg/ml in 50 mM Na-HEPES, pH 8.5 and 30% acetonitrile (v/v) and labelled with respective TMT reagents (Thermo Fisher Scientific, 2.4 mg reagent/1 mg sample) for 1h at RT. Labelling was then quenched with 0.3% hydroxylamine for 15 min at RT and samples acidified (pH^~^2) with formic acid. After verification of labelling efficiency via mass spectrometry, the lysates were mixed in a 1:1 ratio, vacuum dried and desalted on Sep-Pak C18 cartridges.

### Phosphopeptide enrichment

Desalted and vacuum dried samples were solubilized in 1 ml of loading buffer (80% acetonitrile, 5% TFA, 1 M glycolic acid) and mixed with 5 mg of TiO_2_ beads (Titansphere, 5 μm GL Sciences Japan). Samples were incubated for 10 min with agitation, followed by a 1 min 2000 × g spin to pellet the beads. The supernatant was removed and used for a second round of enrichment as explained below. Beads were washed with 150 μl loading buffer followed by two additional washes, the first with 150 μl 80% acetonitrile, 1% TFA and the second with 150 μl 10% acetonitrile, 0.2% TFA. After each wash, beads were pelleted by centrifugation (1 min at 2000 × g) and the supernatant discarded. Beads were dried in a vacuum centrifuge for 30 minutes followed by two elution steps at high pH. For the first elution step, beads were mixed with 100 μl of 1% ammonium hydroxide (v/v) and for the second elution step with 100 μl of 5% ammonium hydroxide (v/v). Each time beads were incubated for 10 min with agitation, and pelleted at 2000 × g for 1 min. The two elutions were removed following each spin, and subsequently pooled together before undergoing vacuum drying. The supernatant from the TiO_2_ enrichment was desalted on Sep-Pak C18 and the High Select Fe-NTA phosphopeptide enrichment kit (Thermo Fisher Scientific) was used according to manufacturer’s instructions for a second round of enrichment.

### Sample fractionation and desalting

Combined TiO_2_ and Fe-NTA phosphopeptide eluates (phosphoproteome) as well as 100 μg of the post-enrichment supernatant (total proteome) were fractionated using the Pierce High pH Reversed-Phase kit (Thermo Fisher Scientific) according to manufacturer’s instructions. Resulting fractions, 8 for each phospho- and total proteome, were taken to dryness by vacuum centrifugation and further desalted on a stage tip using Empore C18 discs (3M). Briefly, each stage tip was packed with one C18 disc, conditioned with 100 μl of 100% methanol, followed by 200 μl of 1% TFA. The sample was loaded in 100 μl of 1% TFA, washed 3 times with 200 μl of 1% TFA and eluted with 50 μl of 50% acetonitrile, 5% TFA. The desalted phosphoproteome and total proteome fractions were vacuum dried in preparation for LC-MS/MS analysis.

### LC-MS/MS and data processing

Samples were resuspended in 0.1% TFA and loaded on a 50 cm Easy Spray PepMap column (75 μm inner diameter, 2 μm particle size, Thermo Fisher Scientific) equipped with an integrated electrospray emitter. Reverse phase chromatography was performed using the RSLC nano U3000 (Thermo Fisher Scientific) with a binary buffer system (solvent A: 0.1% formic acid, 5% DMSO; solvent B: 80% acetonitrile, 0.1% formic acid, 5% DMSO) at a flow rate of 250 nl/minute. The samples were run on a linear gradient of 5-60% B in 150 min with a total run time of 180 min including column conditioning. The nanoLC was coupled to an Orbitrap Fusion Lumos mass spectrometer using an EasySpray nano source (Thermo Fisher Scientific). The Orbitrap Fusion Lumos was operated in data-dependent mode using 3 acquisition methods. Phospho MS2 method: HCD MS/MS scans (R=50,000) were acquired after an MS1 survey scan (R=120, 000) using MS1 target of 4E5 ions, and MS2 target of 2E5 ions. The number of precursor ions selected for fragmentation was determined by the “Top Speed” acquisition algorithm with a cycle time of 3 seconds, and a dynamic exclusion of 60 seconds. The maximum ion injection time utilized for MS2 scans was 86 ms and the HCD collision energy was set at 38. Phospho MS3 method: CID MS/MS scans (R=30,000) were acquired after an MS1 survey scan with parameters as above. The MS2 ion target was set at 5E4 with multistage activation of the neutral loss (H3PO4) enabled. The maximum ion injection time utilized for MS2 scans was 80 ms and the CID collision energy was set at 35. HCD MS3 scan (R=60,000) was performed with synchronous precursor selection enabled to include up to 5 MS2 fragment ions. The ion target was 1E5, maximum ion injection time was 105 ms and the HCD collision energy was set at 65. Proteome MS2 method: same as Phospho MS2 with the following modification to MS2 target 5E4 ions. Acquired raw data files were processed with MaxQuant (44) (version 1.5.2.8) and peptides were identified from the MS/MS spectra searched against *Toxoplasma gondii* (ToxoDB) and *Homo sapiens* (UniProt) proteomes using Andromeda (45) search engine. TMT based experiments in MaxQuant were performed using the ‘reporter ion MS2 or MS3’ built-in quantification algorithm with reporter mass tolerance set to 0.003 Da. Cysteine carbamidomethylation was selected as a fixed modification. Methionine oxidation, acetylation of protein N-terminus, deamidation (NQ) and phosphorylation (S, T, Y) were selected as variable modifications. The enzyme specificity was set to trypsin with a maximum of 2 missed cleavages. The precursor mass tolerance was set to 20 ppm for the first search (used for mass re-calibration) and to 4.5 ppm for the main search. ‘Match between runs’ option was enabled for fractionated samples (time window 0.7 min). The datasets were filtered on posterior error probability to achieve a 1% false discovery rate on protein, peptide and site level. Data were further analyzed as described in the Results section and in the Supplementary Table S1 using Microsoft Office Excel 2016 and Perseus (46) (version 1.5.0.9).

## Data availability

The data behind this manuscript is freely available. The processed proteome and phosphoproteome data is included in Table S1 and we are in the process of placing the raw data on the PRIDE database.

## Acknowledgements

We thank all members of the Treeck laboratory and Dr Ellen Kneupfer for critical discussions. We thank members of the Advanced Imaging and Flow Cytometry Science Technology platforms (STPs) at the Francis Crick Institute for support.

This work was supported by awards to MT by the Francis Crick Institute (https://www.crick.ac.uk/), which receives its core funding from Cancer Research UK (FC001189; https://www.cancerresearchuk.org), the UK Medical Research Council (FC001189; https://www.mrc.ac.uk/) and the Wellcome Trust (FC001189; https://wellcome.ac.uk/). The core facilities at the Francis Crick Institute receive core funding from the MRC, The Wellcome Trust and Cancer Research UK (FC001999). V.B.C was supported by the National Institutes of health grants R01AI120627 (to V.B.C.), and O.L.M by grants T32AI007528 and F31AI118274.

**Figure legends: contained in the main text**

**Supplemental figure legends: in Supplementary figures file**

**Supplemental Material**

Supplementary figures – Fig S1 to S5

Table S1. TvB proteome_phosphoproteome

Supplementary File 1 – Oligonucleotides and synthetic DNA sequences used in this study.

